# Discovery and characterization of unusual O-link glycosylation of IgG4 antibody using LC-MS

**DOI:** 10.1101/2024.04.25.591062

**Authors:** Dariusz J. Janecki, Chi-Ya Kao-Scharf, Andreas Hoffmann

## Abstract

The analysis of several batches of commercial biopharmaceutical product Dupixent using top-down intact mass spectrometry revealed that this immunoglobulin IgG4 features a small amount of O-link glycosylation in Fab region. This is the first report of an O-link glycosylation in IgG4 antibody. The paper describes most likely structure of the O-link glycosylation as well as probable location(s). The relative quantification showed only small quantity of the modification but appearing consistently in several batches of Dupixent. The O-link glycosylation site was characterized by standard tryptic peptide digestion and LC-MS analysis approach.

## 5 INTRODUCTION

Immunoglobulins (Ig) present in human serum contains five primary classes, namely IgG, IgM, IgA, IgD and IgE. IgG represents the most abundant serum immunoglobulin content and serves as the main mediator of humoral immunity (^1^). IgG consists of two heavy and lights chains, connected by disulfide bonds. The protein can be functionally separated into antigen-binding (Fab) and receptor-binding (Fc) region. IgG can be further divided into four subclasses: IgG1, IgG2, IgG3 and IgG4, in the order of decreasing abundance. These four subclasses share overall structure homology but differ slightly in their amino sequence and disulfide bond linkages (^2^).

IgG4 is the least abundant IgG subclass but with unique structural and functional features. It can undergo Fab-arm exchange, which results in monovalent functionality and potential bispecific antigen binding and possess greater propensity for acquiring glycosylation in the variable regions (^3, 4^). Compared to other IgG subclasses, IgG4 has lower affinity to effector molecules, such as Fc receptors and complement. In addition, most of the IgG4 does not activate antibody-dependent immune effector responses. With its’ unique set of properties, IgG4 is often associated with anti-inflammatory effect upon chronic or repeated antigen exposure (^5^).

Glycosylation is one of the most frequent protein post-translational modifications which takes place in the endoplasmic reticulum (^6^). Glycosylation pattern varies across different immunoglobulin classes (^7^). For IgG, all subclasses have one consensus N-linked glycosylation site at asparagine residue within CH2 domain on the Fc region. The majority of glycosylation structures are complex biantennary glycans but lower abundance of high mannose glycans can also be present (^8^). The type of glycan structure has been shown to impact the effector functions of IgG (^9^). N-glycans that lack a core fucose are associated with higher binding to Fc gamma receptor III, which results in enhanced proinflammatory capacity (^9^,^10, 11^). In human IgG, a second glycosylation can occur on the Fab region on the kappa (Vκ), lambda (Vλ) or heavy chain (VH) with no consensus sequence (^8^). Fab glycans are more heavily sialylated and galactosylated than Fc glycans, possibly due to greater accessibility to transferase enzymes. Although not as well investigated as Fc glycosylation, Fab glycosylation can introduce relevant effects on serum IgG (^12^). It is shown that Fab glycosylation impacts the stability, half-life and binding characteristics of antibodies (^13, 14, 15^).

O-linked glycosylation can occur in hinge regions between Fab and Fc portions and has been reported for various immunoglobulins. IgA1 contains nine potential O-glycosylation sites (serine and threonine) in the hinge region, while IgD has been shown to carry four to seven O-glycans (^16, 17, 18^). In addition, partial O-glycosylation has been identified in the hinge region of human IgG3 (^19^). O-linked glycosylation takes place primarily in the Golgi apparatus and its consensus sequence are much less well-defined than N-linked glycans. Regions with high abundance of proline and alanine, along with serine and threonine are well-known sites for O-glycosylation (^7^). O-glycans have different structures from N-glycans but share many of the same monosaccharides. O-glycans are composed of four dominant extensible core structures, with additional sugar such as Gal, *N*-acetylgalactosamine (GlcNAc) and fucose (Fuc). The terminal sugar residues are commonly sialic acids (NeuAc) (^20^). The function of O-glycosylation is not well-understood yet. Several studies indicated that O-glycans in the hinge region of immunoglobulins might prohibit proteolytic degradation. In addition, oligosaccharides in the hinge region can help maintain the protein’s extended conformation, which results in higher flexibility of the Fab region and influence divalent binding to target antigens (^19, 21, 22^).

In the development of biosimilar pharmaceutical product like an antibody the first step is the characterization of the reference molecule (^23,24^). The material can be sourced from different marketed areas and the analyses of these different batches are the foundation of physio-chemical similarity. Dupixent is marketed product (EU, US, JP) containing Dupilumab as an active ingredient (^25^). Dupilumab is a fully human IgG4 monoclonal antibody produced in Chinese Hamster Ovary (CHO) cells by recombinant DNA technology. Dupilumab blocks the action of proteins called interleukins (IL)-4 and IL-13. Both play a major role in causing the signs and symptoms of atopic dermatitis, asthma, chronic rhinosinusitis with nasal polyposis (CRSwNP), prurigo nodularis (PN) and eosinophilic esophagitis (EoE).^25^

Mass spectrometry advances in recent decades have facilitated more robust identification and characterization of glycoproteins and glycopeptides (^26–28^). Newer forms of fragmentation of not only peptide backbone but also glycans themselves allows for gathering more information about glycan structure (^29^). Distinct electro-transfer dissociation (ETD) provides backbone peptide fragmentation with attached intact oligosaccharide (^30,31^). Additionally, the enrichment of glycoprotein using lectins (^32^) or proprietary resins (^33^) produces more material that can be analyzed even when the glycosylation is low abundant. These enrichment techniques can also be quite selective for very specific types of glycosylation (^34–37^).

In this study, we report for the first time of O-glycosylation identified on the Fab region of IgG4. The LC-MS approach with and without enrichment was used successfully to identify and characterize this O-link glycosylation. Moreover, the modification was discovered during characterization of commercially available biologic product Dupixent.

## 6 RESULTS

### 6.1 Intact and subunit analysis by LC-MS

The results of intact mass analysis of deglycosylated with PNGase F Dupixent from EU and US showed additional signals indicating the unexpected PTMs. These additional deconvoluted peaks had masses +660Da and +950Da higher than expected mass of deglycosylated antibody (MMcalc=146899Da). The same samples treated with the FabRICATOR (IdeS) (^38^) enzyme to separate F(ab’)2 and Fc/2 (without deglycosylation) also showed additional signals localized to F(ab’)2 region with similar additional masses. The 2D deconvolution using Genedata workflow is shown in Figure 1 and Figure 2 in Supplementary Information. Although signal intensities were not very strong these results were investigated further.

**Figure 1.**
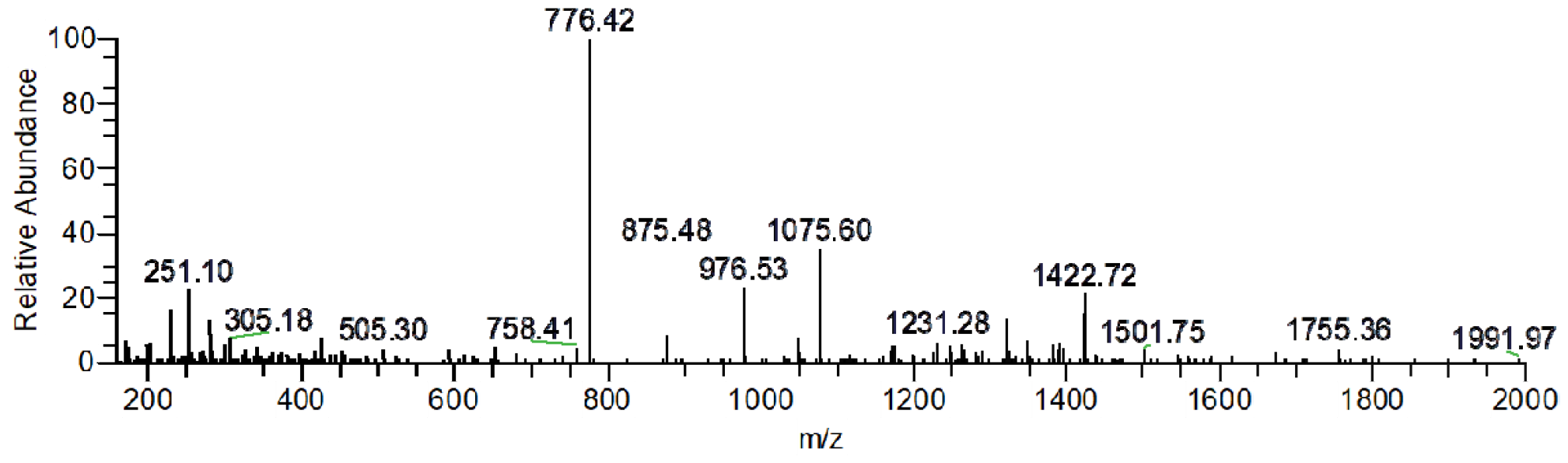
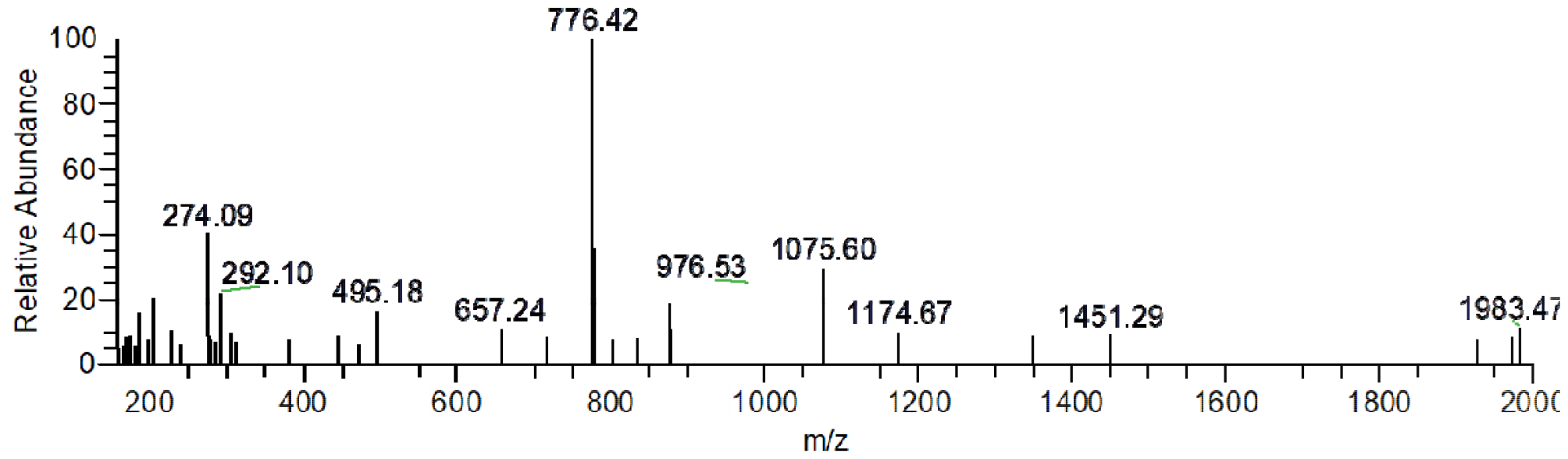
Tandem mass spectra (HCD) of unmodified (top) and modified (bottom) peptide T-H17.

**Figure 2.**
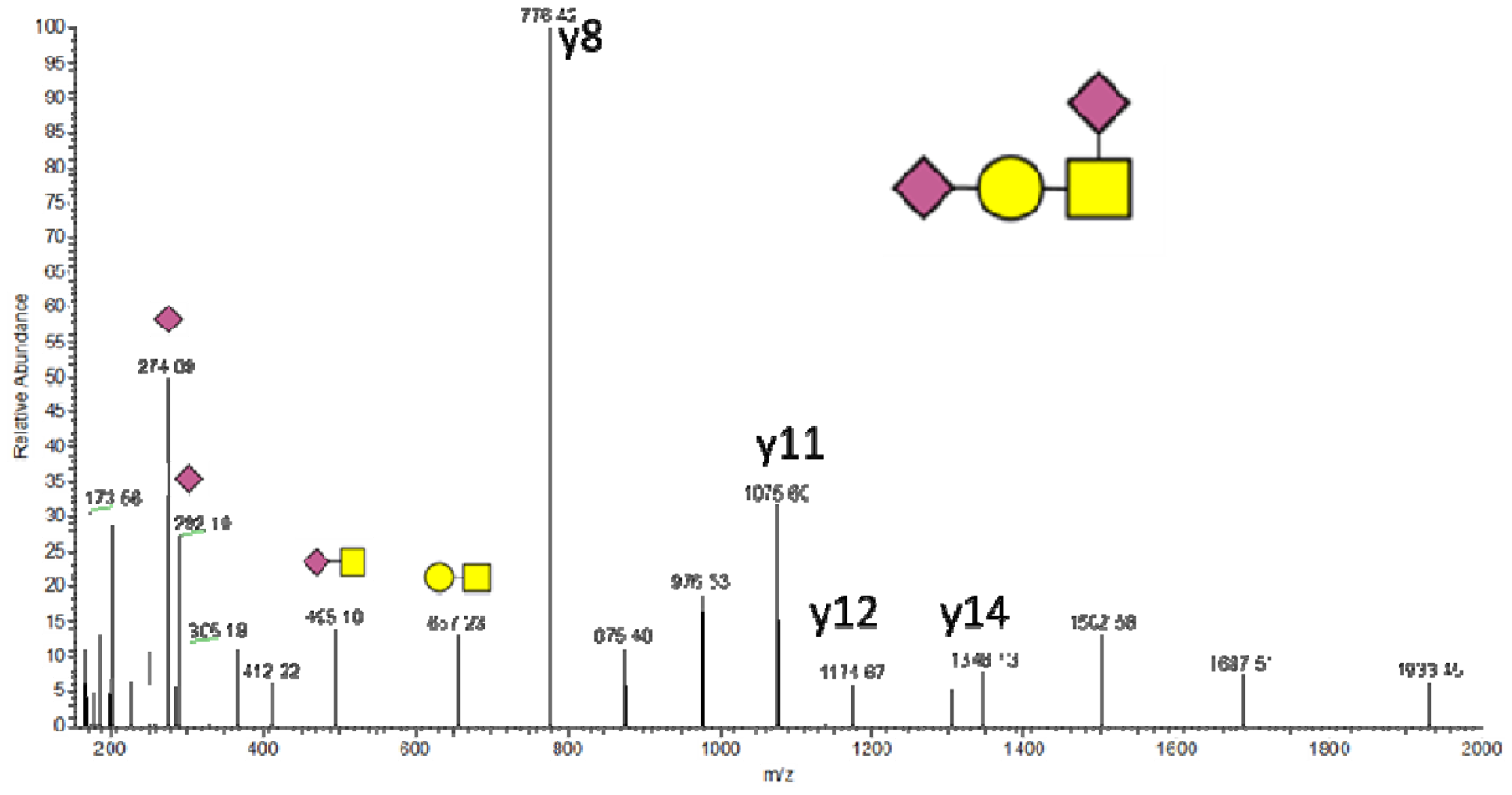
Structure elucidation of the proposed O-link glycan on T-H17 peptide (HCD fragmentation).

The LC-MS analysis of tryptic digest of the samples (data dependent MS/MS using HCD fragmentation) confirmed the presence of additional modifications of heavy chain tryptic peptide T-H17 HC[156-204] DYFPEPVTVSWNSGALTSGVHTFPAVLQSSGLYSLSSVVTVPSSSLGTK. Figure 1 shows HCD fragmentation spectra of unmodified and modified peptide. The fragmentation of ions for unmodified peptide at m/z 1264.89 (+4) show very strong signal for y8 ions at m/z 776.42 Da. These ions are formed from fragmentation of peptide bond N-terminal to proline. The observed mass of unmodified peptide T-H17 was calculated 5055.5450 Da which is in excellent agreement with theoretical mass 5055.5393 Da (error 1 ppm). The lower panel in Figure 1 shows fragmentation of ions at m/z 1501.73 (+4). This spectrum also shows 776.42 Da ions but additionally some ions characteristic for glycosylation (oxonium ions) are present at lower m/z range. These ions are 204.0866, 274.0925, 292.1032, 495.1821 and 657.2330 Da. The mass of the modified peptide was calculated as 6002.8760 Da. The mass difference of the modification can be calculated as 947.3367 Da which points to Hex(1)HexNAc(1)NeuAc(2) O-linked glycosylation. The calculated mass is within 15ppm of expected theoretical mass (947.3230 Da).

The fragmentation was not adequate to place the modification in the peptide. Overall, there are 16 possible places for singly O-link glycosylation (5 Thr and 11 Ser). Although the placement of the modification is difficult, the presence of some characteristic oxonium ions e.g. 495.1821 Da provides some information about the structure of the O-link glycan (^29^). The proposed structure is shown in Figure 2.

The peptide T-H17 was also found with Hex(1)HexNAc(1)NeuAc(1) O-linked glycosylation with the observed mass 5711.7709 Da (1 ppm error). The mass difference if +656.2316 Da correlates to the difference observed in intact mass analysis. The extracted ion chromatogram (see Supplementary Information Figure 3) suggests that there are at least two forms of the peptide -possibly with different positioning of the NeuAc or different position of the whole O-link glycan within the peptide. Several other fragments (possibly clipping) of the T-H17 peptide with O-link were also detected.

**Figure 3.**
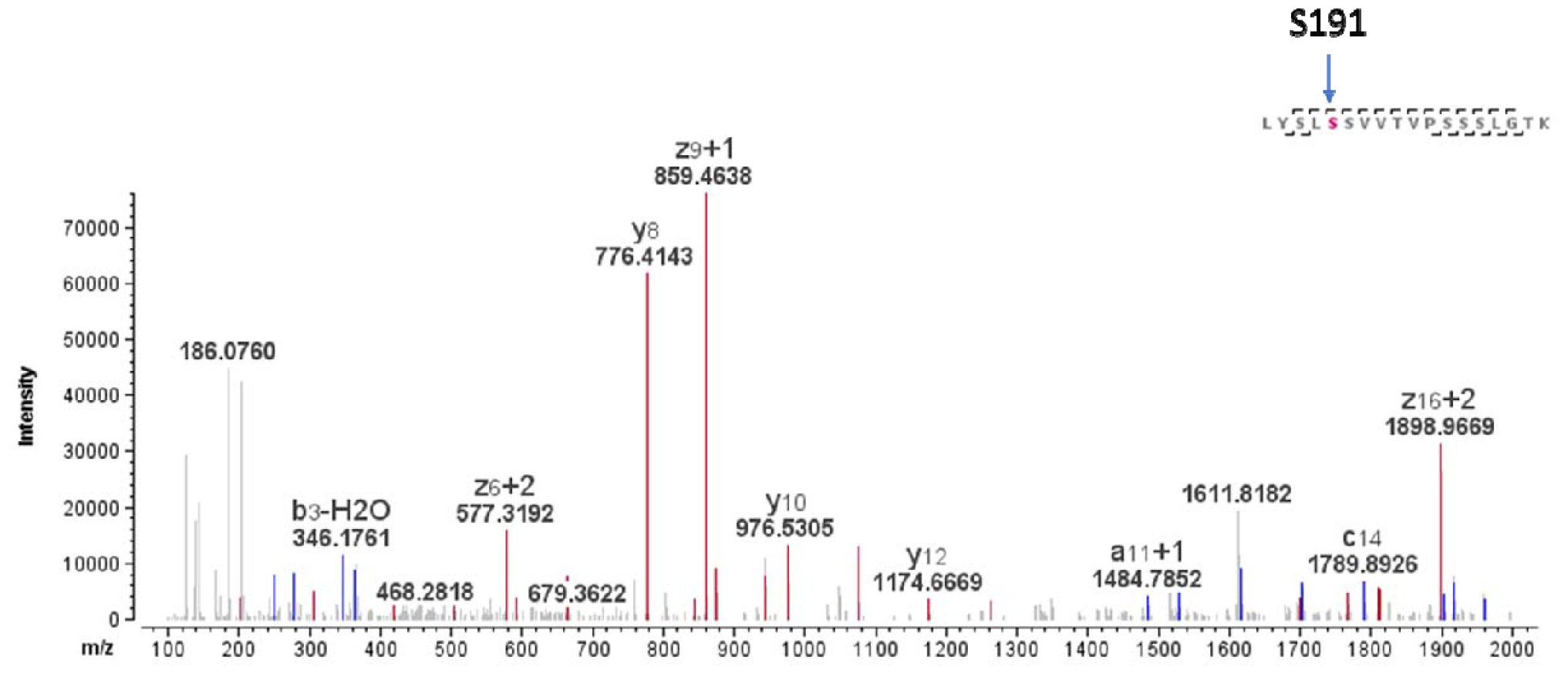
EThcD fragmentation mass spectrum: the position of S191 occupied by O-link glycosylation.

While HCD fragmentation can be helpful with glycan structures it can provide little information about the placement of the modification. For this reason, the data dependent acquisition with electron transfer dissociation (ETD) or electron-transfer/higher-energy collision dissociation (EThcD) fragmentation modes was employed.

The sample for ETD/EThcD experiments was enriched first using GLYCOCatch column from Genovis. The sample for enrichment was Dupixent batch from EU market. The GLYCOCatch protocol employs the use of SialEXO enzyme which removes terminal sialic acid from O-link glycans. The final enriched protein was reduced and alkylated and digested with Lys-C. The expected modification after sialic acid removal was Hex(1)HexNAc(1) at the site of the glycosylation.

The examples of EThcD spectra with best assignment of the glycosylation positions are presented in Figure 3 and Figure 4. Please note that the peptides observed are not full T-H17 peptide but rather fragments of it. It was observed in previous digestions (tryptic) that some additional clipping was occurring in that region of the mAb. This was also observed in intact mass data (data not shown).

**Figure 4.**
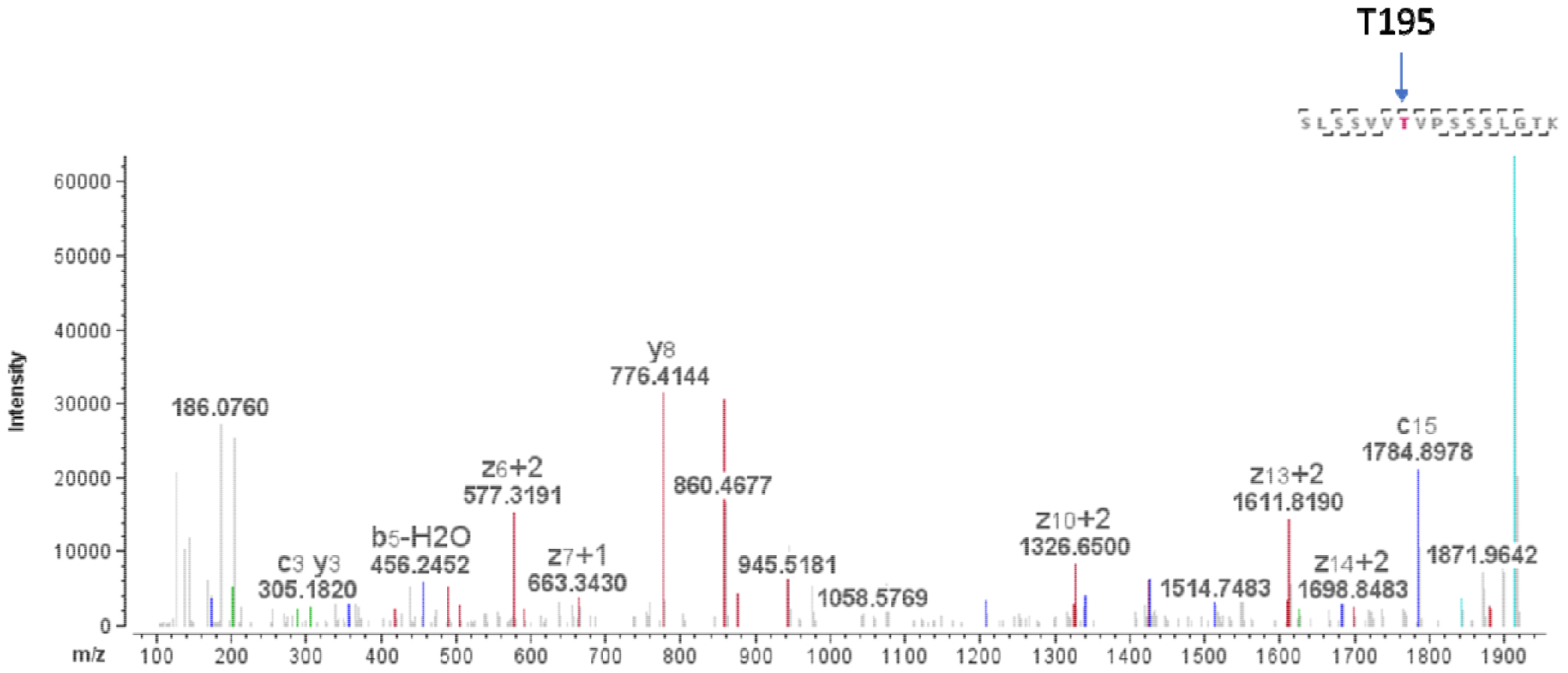
EThcD fragmentation mass spectrum: the position of T195 occupied by O-link glycosylation.

During careful data analysis several position could be assigned with high probability. Namely S191, S192, T195 and S198. Unfortunately, not all EThcD spectra were pointing clearly at specific Ser or Thr and it is very likely that numerous positions are occupied. Double O-link glycosylations were also observed but only with mass. See Table 1 for results of EThcD fragmentation and Table 2 for MS only matches of intact T-H17 peptide.

**Table 1.**
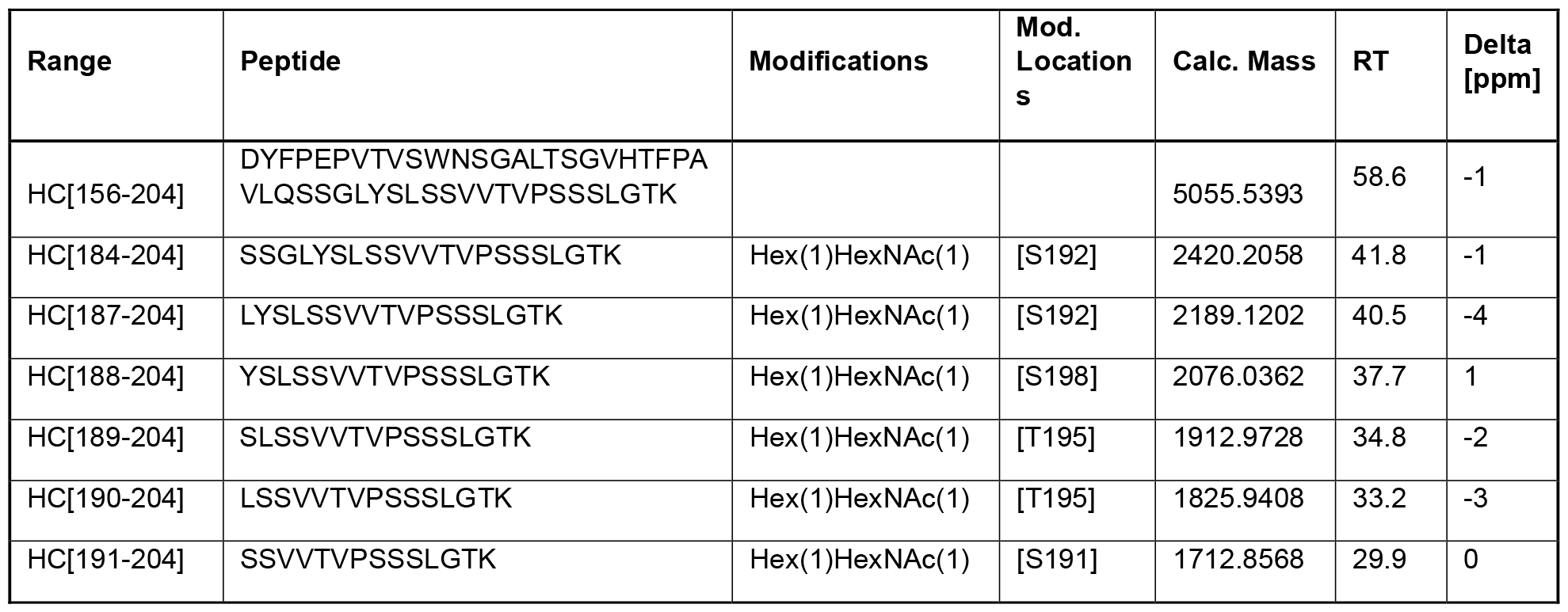
O-link modifications detected with MS/MS data (GlycoCATCH enriched sample)

**Table 2.**
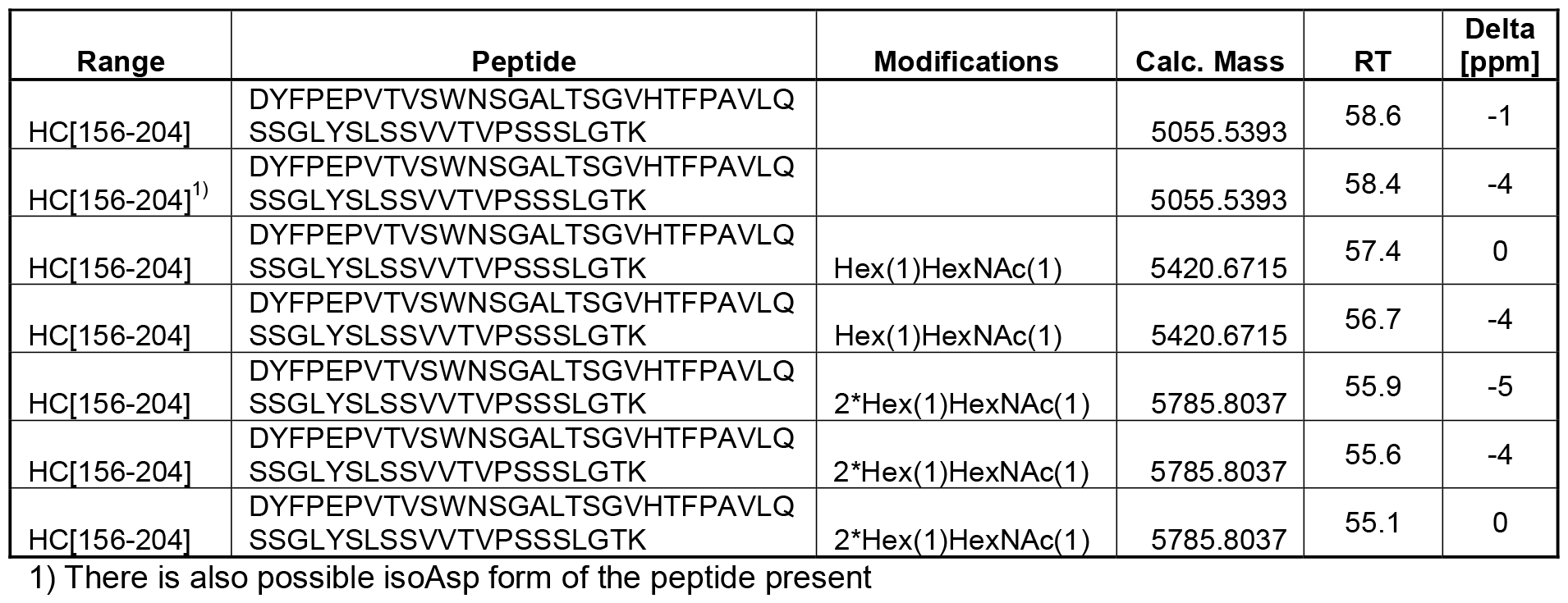
Full length peptide with O-link glycosylation matches by MS only (GlycoCATCH enriched sample).

Targeted LC-MS approach with ETD fragmentation did not provide additional information. Despite strong signal of the precursor ions selected in data dependent approach, the fragmentation was not always optimal, and these additional LC-MS experiments did not add any further modification site(s) information (data not shown).

## 7 DISSCUSSION

Multiple post-translational modifications play a critical role in defining the development target of a biosimilar, requiring thorough characterization of the reference product. Mass spectrometry-based analytics showed an unusual, low abundant modification in an IgG4. The intact and subunit data revealed additional mass signals around +660 Da and +950 Da suggesting the presence of O-link glycosylation localized in F(ab’)2 region. The O-link glycosylation is rare in monoclonal antibodies and only described in some isotypes (^39^) e.g. IgG3, IgA. Thus far IgG4 isotype has not been described to contain O-link glycosylation sites. The identified O-link glycosylated peptide HC[156-204] DYFPEPVTVSWNSGALTSGVHTFPAVLQSSGLYSLSSVVTVPSSSLGTK is common in all IgG4 antibodies and highly conserved in IgG1. Comparison of available X-ray structures of several IgG4, including Dupilumab, revealed high structural similarity of the peptide region of the identified O-glycan sites. Thus, we analyzed two other marketed biologic products that contain an IgG4 antibody as active substance (Nivolumab, Pembrolizumab) for similar O-glycans. However, no O-link glycosylation was found in these products (data not shown). Also, several different IgG1 biologic products revealed no O-linked glycosylation (data not shown). This finding is in line with the observation that O-linked glycans predominantly prevail in disordered regions. While present in the extended hinge of IgG3 (^19^), such regions are not present in IgG4 or IgG1. However, some O-glycosylation sites are found in or close to loops or turn motifs of structural domains as observed for Dupilumab. The reason for Dupilumab’s unique feature is unclear. It may be related to its specific Fab region impacting the transport through endoplasmic reticulum and Golgi apparatus, thereby impacting the glycosylation pattern.

Several other batches of Dupixent from EU and US sources were also analyzed using the reduced tryptic peptide mapping approach and all batches have detectable levels of the O-link glycosylation on the T-H17 peptide. The relative quantification of these modifications is summarized in Table 3. The O-link glycosylated T-H17 peptide with one sialic acid was eluting at two retention times in our LC gradient and each glycosylation was quantified separately. The difference in the hydrophobicity of the species can be due to different position of sialic acid in O-link glycan (see Figure 2) but more likely different position of the O-link glycosylation within backbone of the T-H17 peptide. It was shown that at least four positions (three Ser and one Thr) are likely occupied with the O-link glycosylation in this peptide (see Table 1 for details).

**Table 3.** Relative quantification of O-link glycosylation of T-H17 peptide of different batches of Dupixent.

| Name | EU Batch1 | EU Batch2 | EU Batch3 | EU Batch4 | EU Batch5 | EU Batch6 | EU Batch7 | US Batch1 | US Batch2 | US Batch3 | US Batch4 |
| --- | --- | --- | --- | --- | --- | --- | --- | --- | --- | --- | --- |
| T-H17 O-link SA1 | n.d. | n.d. | 0.04 | 0.04 | 0.01 | 0.02 | 0.02 | n.d. | 0.02 | 0.02 | 0.02 |
| T-H17 O-link SA1 | 0.01 | 0.01 | 0.02 | 0.01 | 0.01 | 0.02 | 0.02 | 0.01 | 0.02 | 0.02 | 0.02 |
| T-H17 O-link SA2 | 0.06 | 0.08 | 0.02 | 0.02 | 0.02 | 0.03 | 0.02 | 0.09 | 0.02 | 0.02 | 0.02 |

n.d. – not detected

The relevance of the observed O-link glycosylation for biosimilar development is limited. Due to its low abundance no impact on safety and efficacy is expected. Nevertheless, it shows the importance to perform an unbiased initial characterization of the reference product to avoid missing potentially important quality attributes when setting development targets for the biosimilar. Another low abundance modification in the T-H17 peptide was N-link glycosylation at the reverse consensus site at N167 asparagine. Occupation of similar sites was described in literature (^40^).

## 8 MATERIALS and METHODS

### 8.1 CEX fractionation of Dupixent

The samples were fractionated by CEX method. Dupixent (prefilled syringe 300 mg/2mL) from EU (c=175.3 mg/mL) and US (c= 148.8 mg/mL) were diluted to about 30 mg/mL so they are not too viscous for the HPLC system. An aliquot of 250 μL was injected to the HPLC system (Agilent 1260). The collected fractions were concentrated and re-buffered using the buffer consisting of 10mM succinic acid, 150 mM proline and 50 mM arginine hydrochloride, pH 5.9. Concentration of fractions was achieved by using Vivaspin 20 ultrafiltration units in combination with an Eppendorf 5810R centrifuge (3900 rpm, 4°C). Each fraction was further re-buffered to a final volume of 1000 μL. Samples were frozen at -20°C before further analysis.

### 8.2 Intact and subunit mass analysis

The CEX fractions were diluted with DPBS to concentration about 1 mg/mL. An aliquot of 50 μL was deglycosylated by adding 1.5 μL of PNGase F enzyme (New England Labs, P0710S) and incubated for 30 min at 50°C on thermomixer (without shaking). Another 50 μL aliquot was treated with FabRICATOR (IdeS) enzyme (Genovis, A0-FR1-096). A tube of enzyme was reconstituted with 100 μL DPBS and 50 μL of the enzyme was mixed with 50 μL of the sample. The vials were incubated in thermomixer for 60 min at 37°C @350rpm.

### 8.3 Reduced tryptic digestion

The samples to be digested were mixed with freshly prepared denaturation buffer (7.5M Guanidine HCl, 100 mM Tris HCl, pH 8.3) and DTT and incubated for 30 min at 55°C. (Guanidine HCl, Sigma, part# 50937; Tris HCl, Merck, part # 1.08386.

In general, 46.8 μL of denaturing buffer was mixed with 1.8 μL of 0.5 M dithiotreitol DTT (Sigma, part # 43815) and protein solution corresponding to 200 μg of protein. Additionally, 65.6 μL miliQ water is added. After reduction with DTT the samples are cooled down and 3.4 μL of iodoacetic acid (IAAc, 0.5M, freshly prepared, Sigma, part # 57857) was added. The alkylation reaction was performed at room temperature in darkness for 15 minutes. To quench the reaction 1.0 μL of 0.5M DTT was added. Before the digestion, a buffer exchange was performed using BioSpin 6 column (BioRad, part# 732-6222) following the manufacturer’s protocol. The final reduced and alkylated flow through from the column was mixed with 1 mg/mL solution of trypsin enzyme (Promega, V511A) in ratio 1:10 (w/w) enzyme to protein. The digestion was performed for 2 hours at 37°C in a thermomixer (Eppendorf) without shaking. To stop the digestion 1 μL of 10% trifluoroacetic acid (TFA) was added.

### 8.4 Enrichment of O-link glycosylated antibody

The method was used as described in Genovis GlycoCATCH enzyme protocol (^33^). After the sample was enriched a Lys-C digestion was performed. 100 μL of the sample (Dupixent EU, c=153.8 mg/mL) was mixed with 100 μL of DPBS (Thermo Scientific, (J67802.AP)) as binding buffer. SialExo enzyme that is part of the GlycoCATCH kit (GlycoCATCH Affinity Purification; Genovis (G3-OC6-002)) was reconstructed in 20 μL miliQ water to final concentration 10 units/μL. The GlycoCATCH column was washed 3 times with 300 μL of binding buffer (DPBS) centrifuged down for 1 min at 1000 g each time. After the column was washed the sample in DPBS was pipetted onto the column. Additionally, 10μL of SialEXO solution (50 units) was added in and the column was slowly mixed for 2 h at room temperature on up-down shaker. The column was placed in Eppendorf tube and centrifuged for 1 min at 1000 g. The flow through was not analyzed. Next, the column was washed seven times according to procedure with 0.4 M NaCl (NaCl (EMD Millipore; 1.06404.1000) in DPBS. After final centrifuge a 50 μL aliquot of Elute Buffer (8 M urea; Urea (Sigma, U6504-500G) in DPBS) was added. The column was equilibrated for 5 min and then centrifuged for 2 min at 1000 g and the elute collect to a new Eppendorf tube. The elution was repeated second time to obtain ∼100 μL of elute.

#### 8.4.1 Lys-C Digest of the enriched sample

The GlycoCATCH enriched sample was reduced and alkylated and subsequently digested with Lys-C enzyme. The sample was reduced with 3 μL of freshly prepared 1 M DTT (Sigma, #43815-5G) for 15 min at 60°C, and after cool down alkylated with 6 μL of freshly prepared 1 M iodoacetamide (IAA, Sigma, #I1149-5G). The alkylation was performed for 15 min at room temperature in darkness. To quench the alkylation additional 2 μL of DTT was added. In order to lower the concentration of urea the reduced and alkylated sample was first diluted with 300 μL miliQ water and 100 μL of 1M Tris pH 8.0 (Life Technologies, 15568025), (final concentration of urea 1.6 M and pH around 8.0). The digestion was performed by adding 20 μL of 0.5 mg/mL Lys-C enzyme (Waco FujiFilm, 125-05061) and incubating for 5 h at 37°C. The enzyme reaction was quenched with 10 μL of 10% TFA (Thermo; P#85183).

### 8.5 LC-MS analysis

#### 8.5.1 Intact and subunit mass analysis

LC-MS analysis was performed using Thermo QExactive Plus mass spectrometer attached to Agilent LC system (1260). The column was BioResolve RP mAb Polyphenyl (Waters, 186008946) with eluent A being 0.08% TFA, 0.02% formic acid (FA) in MiliQ water and eluent B was 0.08% TFA, 0.02% FA in ACN. The column temperature was 60°C and the gradient was from 27% B to 42% B over 15 min with the flow of 0.5 mL/min. The QExactive Plus H-ESI source was in positive mode, Spray Voltage 3800V, In-source CID 40eV, resolution 35,000; 10 microscans, Protein Mode ON. These settings were applied to intact deglycosylated sample as well as second Segment (eluting F(ab’)2) of IdeS digested sample.

#### 8.5.2 Peptide mapping

LC-MS peptide mapping of the sample was performed using Agilent LC (for tryptic peptide mapping) or Vanquish LC (for Lys-C digested enriched sample). The HPLC systems were coupled to an Orbitrap Fusion Tribrid mass spectrometer (Thermo Fisher Scientific). Reversed phase (RP) chromatography was performed using a Acquity UPLC Peptide BEH C18 column (Waters, 186003687), 1.7 μm particle size, 2.1 × 150 mm at a flow rate of 300 μL/min and 60 °C column temperature. Solvents contained MiliQ water and 0.1% TFA (solvent A) as well as 0.1% TFA in 100% acetonitrile (Merck, 1.00030) (solvent B). 20 μg of the digest was injected onto the column and eluted using a gradient from 0 to 70 min with increasing solvent B from 0 to 44%. For the tryptic digest the acquisition was performed with the following parameters: H-ESI ion source (positive ion, 3500 V); Orbitrap resolution (120,000 in MS and 30,000 in MS2); mass scan range: 300–2000 m/z; RF lens: 60%; standard AGC target; maximum injection time: 60 ms; 1 microscan; data dependent MS2 acquisition with HCD activation and stepped Collision Energy Mode (20,35,50%).

#### 8.5.3 ETD and EThD fragmentation

The same LC gradient was used as for tryptic peptide mapping experiment, but the MS settings included data dependent ETD fragmentation or ETD followed by additional HCD activation (EThD). This first approach was not targeted. Additionally, the second approach was to create inclusion list with possible T-H17 peptide O-link modifications. The same LC conditions were used for this approach.

### 8.6 Data analysis

The Genedata Expressionist® 16.5.5 Refiner MS (Genedata) package was used. For the analysis of Intact deglycosylated samples Raw files were loaded into Load from File activity and processed with the following activities and respective settings: Intact Protein (10–200 kDa, mass step 0.1 Da, Maximum Entropy Deconvolution, Protonation), Manual Peak Edit, (manually edit the 2D peaks of interest); and Export Excel and PDF Reports. For analysis of IdeS digested intact samples additional Selection (selecting the segment of data with F(ab’)2 peak) activity was added since the data acquisition on QE plus was in two segments.

Tryptic peptide mapping was performed with Genedata workflows with MS Library constructed for this purpose (see Table 1 in Supplemental Information). The analysis of Lys-C digestion of enriched samples was performed using Genedata general peptide mapping activity with settings: Mass Tolerance = 10ppm; m/z Mass Tolerance = 10ppm; Enzyme = SemiTrypsin; Min Peptide Length = 3; Fixed Modification = Carbamidomethyl (C); Variable Modification = Hex(1)HexNAc(1) (ST).

## Notes

### Competing Interest Statement

The authors have declared no competing interest.

