## Supplemental Information for "Discovery and characterization of unusual O-link glycosylation of IgG4 antibody using LC-MS"

### 1 Supplemental information

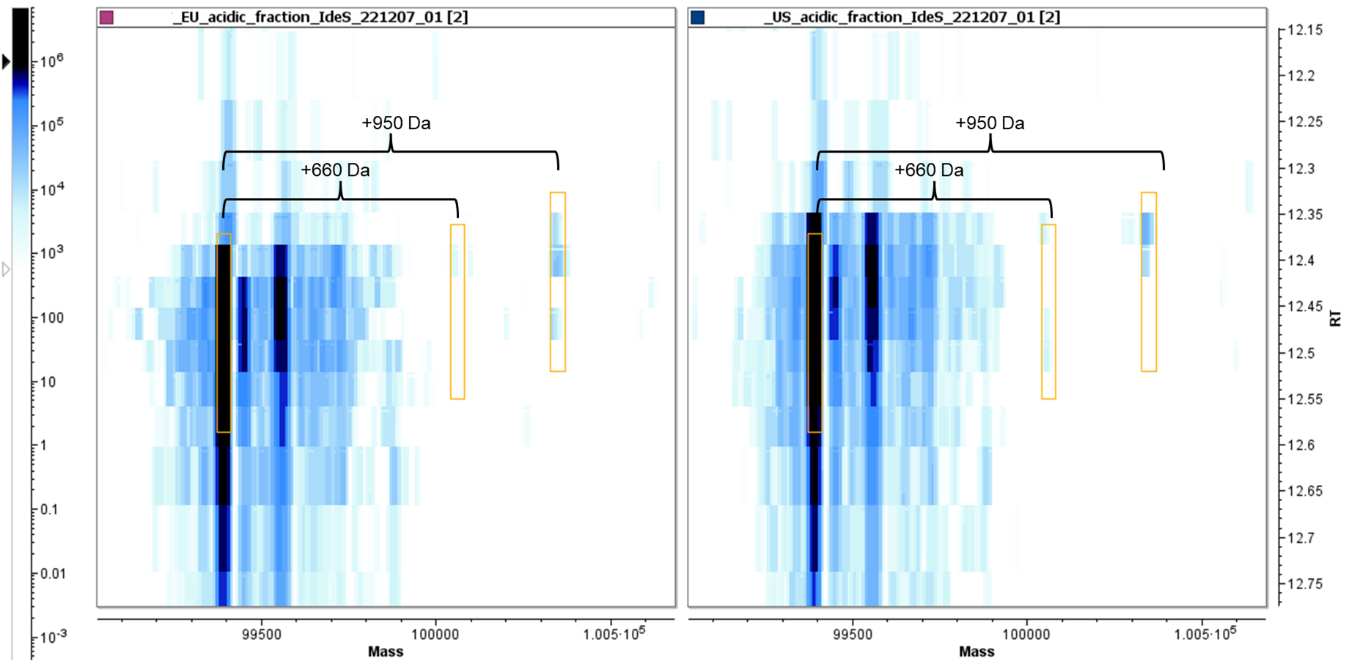
Figure 1 2D mass deconvolution of IdeS digest (F(ab’)2) of Dupixent EU and US acidic fractions

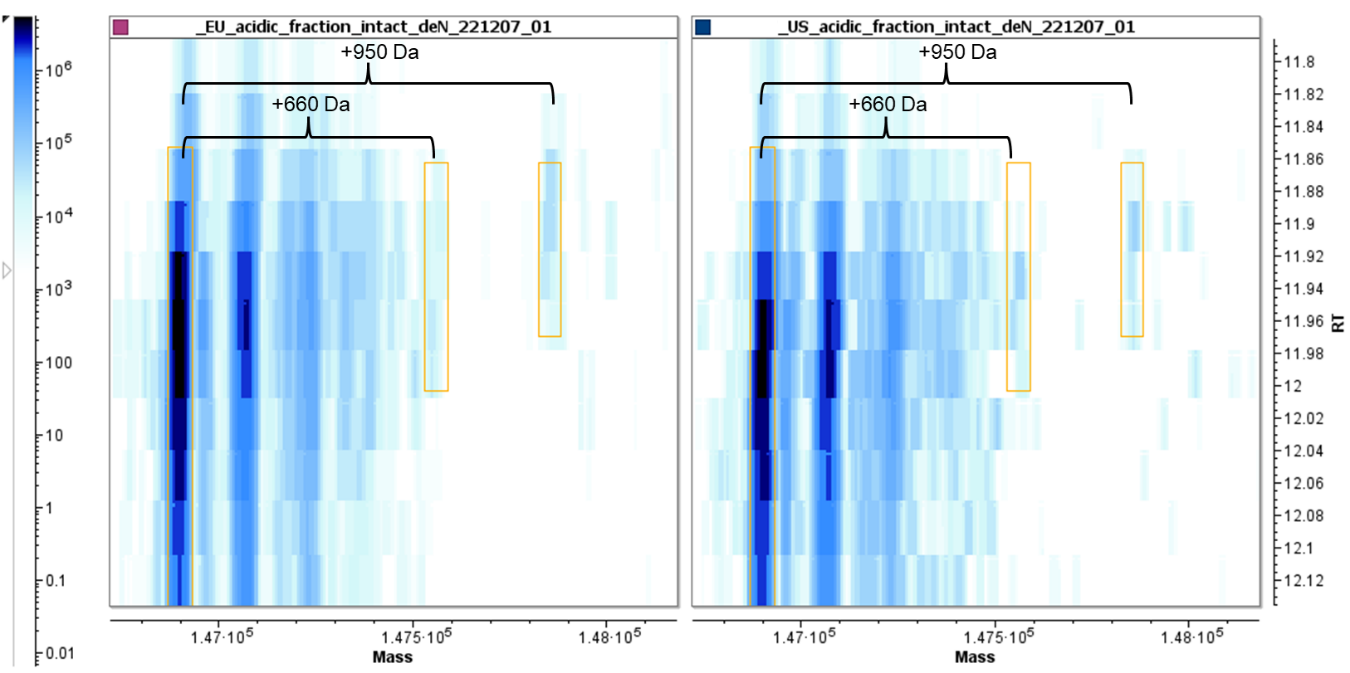
Figure 2 2D intact mass deconvolution of PNGase F treated of Dupixent EU and US acidic fractions

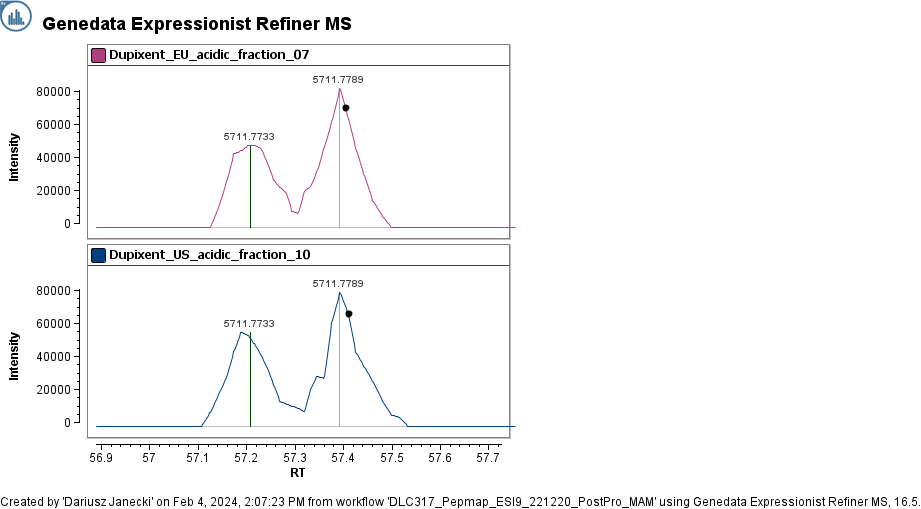
Figure 3 Extracted Ion Chromatogram for O-link glycosylation with one sialic acid on T-H17 peptide

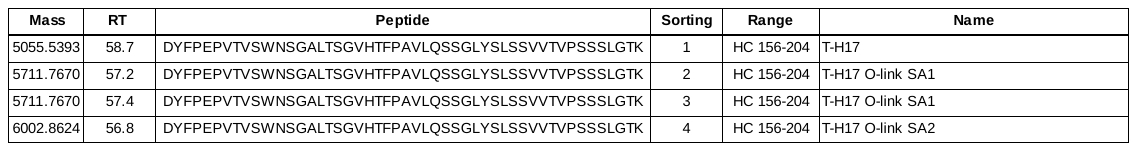
Table 1 MS Library used for Genedata peptide mapping workflow
